# A Drug Screening Platform for Protein Expression Levels in Neurological Disorders

**DOI:** 10.1101/2024.02.27.582434

**Authors:** Farida Emran, Ibrahim Kays, Chiu-An Lo, Yueyang Li, Brian E. Chen

## Abstract

Neurological and psychiatric diseases and disorders affect more than half of the population. Many of these diseases are caused by the malfunctioning of protein synthesis, where too little or too much production of a protein harms a cell and its functions within the brain. We developed a drug screening platform to identify compounds that target the primary cause of these diseases, namely protein expression amounts. This cellular assay monitors protein expression of a target disease gene along with the protein expression of a control gene using the Protein Quantitation Ratioing (PQR) technique. PQR tracks protein concentration using fluorescence. We used human cells and CRISPR-Cas9 genome editing to insert the *Protein Quantitation Reporter* into target genes. These cells are used in high-throughput drug screening measuring the fluorescence as the assay. Drug hits can be validated using the same PQR technique or animal models of the disease.

**Highlights:**

1. The assay can identify drugs that directly address the molecular cause of a disease.
2. The Protein Quantitation Ratioing (PQR) technique allows for tracking and measuring protein amounts over time in single living cells before, during, and after drug administration.
3. Genome editing to insert the *PQR* into the target gene allows tracking of endogenous protein expression.
4. Using human cell lines allow for faster production of knock-in cells.
5. Patient mutations can be replicated using genome editing during the knock-in step.
6. Using induced pluripotent stem cells allow for an unlimited supply of genome edited differentiated cells such as neurons with the *PQR* knock-in reporter.

## Introduction

Protein synthesis within cells is a tightly regulated process where too little or too much of a single protein can cause diseases and disorders such as cancer, neurodegenerative diseases, and psychiatric disorders. Despite having two copies of the genome, there are more than 1,000 disorders and diseases caused by loss of only one copy of a gene, called haploinsufficiency, where the expression of the remaining gene is insufficient to prevent the disorder. More than 20% of haploinsufficiencies result in neurological disorders (Dang et al., 2008; Huang et al., 2010).

Tracking protein expression over time in cells is a useful tool to interrogate the mechanisms of these genetic diseases. The Protein Quantitation Ratioing (PQR) technique measures protein production of any gene over time in single cells *in vivo* (Lo et al., 2015). The PQR technique works by inserting the *Protein Quantitation Reporter* DNA sequence into any gene of interest. During protein synthesis, the PQR peptide sequence produces one molecule of a fluorescent protein reporter (e.g., GFP) for every one molecule of protein of interest (**Figure 1a**). The GFP molecule is produced at the protein translation step and is separated from the protein of interest, and so does not interfere with the endogenous protein’s activity. Because the GFP is produced in a 1:1 equimolar ratio with the protein of interest, the GFP fluorescence intensity (i.e., brightness) of the cell is used as a readout for protein abundance.

**Figure 1.**
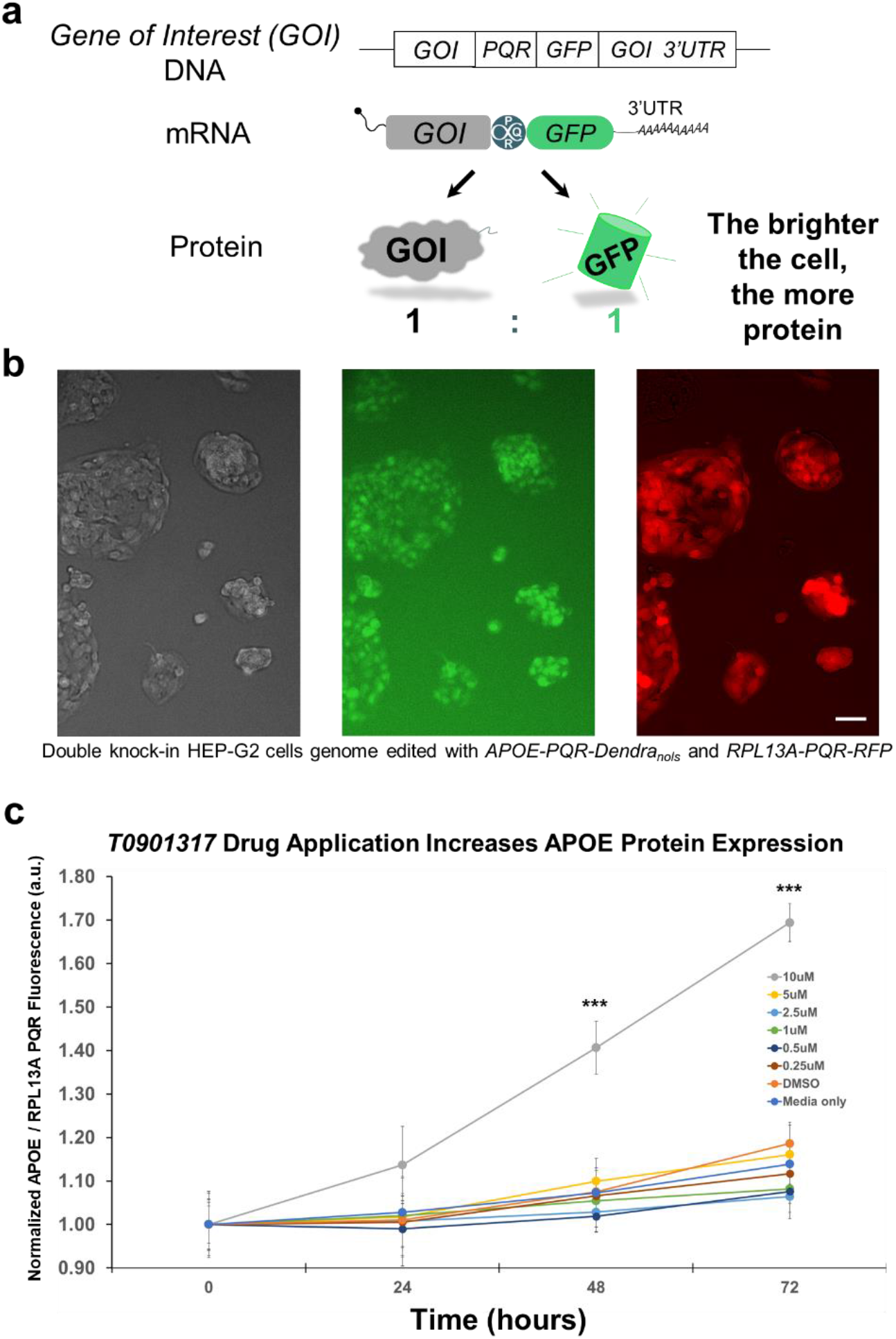
Protein Quantitation Ratioing (PQR) can measure endogenous protein expression over time in single living cells. (a) Knock-in of a *Protein Quantitation Reporter* (*PQR*) into a gene of interest (GOI) using genome editing can quantify the protein levels using the amount of GFP expressed in a cell (i.e., brightness). The *PQR* creates a polycistronic mRNA for co-transcription and co-translation of GFP and the gene of interest. The *PQR* construct allows for one molecule of GFP to be synthesized for every one molecule of the protein of interest synthesized. The fluorescence intensity of a cell can then be used to quantitate its concentration. (b) Representative images of human HEP-G2 hepatocytes expressing APOE-PQR-Dendra_nols_ and RPL13A-PQR-RFP. The brightness of the Dendra in the green channel quantifies the steady state amount of APOE for each cell, and in the red channel, the steady state amount of RPL13A protein for each cell. Scale bar is 25 µm. (c) 10µM T0901317 significantly increased endogenous APOE protein expression after two days in HEP-G2 cells (*p* < 0.001). APOE fluorescence in the green channel was normalized to the control RPL13A protein amounts in the red channel, expressed in arbitrary units (a.u.). Error bars are standard deviation.

Using CRISPR-Cas9 genome editing, the *PQR* can be inserted into any gene of interest to report endogenous protein production, including in *Drosophila melanogaster* (Lo and Chen, 2019), zebrafish (Armstrong et al., 2016), mouse (Bagheri et al., 2018; Kays and Chen, 2019), and human genomes (Kays and Chen, 2019; Lo et al., 2017; Lo et al., 2015). For example, we have used PQR to compare mRNA versus protein amounts in single cells (Kays and Chen, 2019), or to measure the protein expression dynamics for both parental alleles simultaneously in cells over time in the living animal (Lo and Chen, 2019). Here, we demonstrate the use of PQR to quantify protein expression of disease-associated genes in human cells. These cells are then used in high-throughput drug screens based on their PQR fluorescence to identify compounds that can increase the target gene’s protein amounts.

## Results

### T0901317 Drug Application Increases APOE Protein Expression

We first sought to verify that PQR can measure drug-induced changes in protein amounts in human cells over time. We chose the drug T0901317, which increases Apolipoprotein E (APOE) protein and mRNA via activation of liver X receptor alpha (Beyea et al., 2007; Kurano et al., 2011; Liang et al., 2004; Riddell et al., 2007). We used CRISPR-Cas9 genome editing to insert a *PQR* into the *Apolipoprotein E* gene, whose variants are the most common genetic risk factor for Alzheimer’s disease. We generated *APOE-PQR-Dendra2*_*nols*_ and *RPL13A-PQR-RFP*, double knock-in human HEP-G2 cells (**Figure 1b**). Dendra2 is a photoconvertible green to red fluorophore (Chudakov et al., 2007), and the nucleolar localization signal (nols) is used to sequester the fluorophore into the nucleolus for ease of imaging and analysis (Lo and Chen, 2019; Lo et al., 2015). We use the *RPL13A* gene as a control (Kays and Chen, 2019; Lo and Chen, 2019; Lo et al., 2017; Lo et al., 2015) in a second imaging channel for changes in imaging conditions, imaging artifacts from the plate, cell health, and drugs that non-selectively alter protein expression rather than just the target gene.

Previous studies have used concentrations of T0901317 ranging from 2nM and 10µM over timescales of 24 hours to 7 days in different cell types ranging from human THP-1 macrophage-like cell lines to HEP-G2 cell lines to mice *in vivo* (Beyea et al., 2007; Joseph et al., 2002; Kurano et al., 2011; Liang et al., 2004; Quinet et al., 2004; Riddell et al., 2007; Whitney et al., 2002). After T0901317 administration, *APOE* mRNA levels increased between zero to 3-fold, and APOE protein increased between zero to 2-fold, depending on the cell type and conditions. The wide range of these previous results is likely due to the differences in cell types used in the studies, and the differences in liver X receptor alpha expression and activation.

Using the *APOE-PQR-Dendra2*_*nols*_; *RPL13A-PQR-RFP* double knock-in human HEP-G2 cells, we imaged APOE protein levels over 3 days in multi-well plates using a high-throughput imager with liquid handling. Application of 10µM T0901317 significantly increased APOE protein by 40% after 2 days and 60% on the next day, without affecting the control protein RPL13A (*p* < 0.001, *n* > 100 cells per well, 28 wells, *t*-test) (**Figure 1c**). Concentrations lower than 10µM did not significantly alter APOE nor RPL13A protein levels.

### High-Throughput Drug Screening for Protein Expression Changes

Using CRISPR-Cas9 genome editing, we inserted a *PQR* into four other genes, *EHMT1, GBA1, SHANK3*, and *SLC6A15* in HEK293T human cells. *EHMT1* is a haploinsufficient gene that causes Kleefstra syndrome, a complex disorder of developmental delay and intellectual disability (Kleefstra et al., 2009). *GBA1* is the most frequent genetic risk factor for Parkinson’s disease (Migdalska-Richards and Schapira, 2016; Murphy et al., 2014), and in many of these cases there is not enough of the GBA1 protein being made (Gegg et al., 2012; Kurzawa-Akanbi et al., 2012; Mazzulli et al., 2011; Murphy et al., 2014). *SHANK3* is a haploinsufficient gene that causes Phelan-McDermid syndrome but whose changes in protein levels is are also associated with autism and obsessive-compulsive disorder (Yi et al., 2016). *SLC6A15* is a gene implicated in stress susceptibility and depression (Chandra et al., 2017). We generated *EHMT1-PQR-GFP, GBA1-PQR-GFP*_*nols*_, *SHANK3-PQR-RFP*_*nols*_, or *SLC6A15-PQR-RFP*_*nols*_ HEK293T cells (**Figure 2a**). We also demonstrated that patient mutations can be replicated during the *PQR* insertion genome editing step. For example, we recreated the *GBA1* patient mutation L444P (**Figure 2a**) that is one of the most common mutations in the GBA1 protein (Cilia et al., 2016).

**Figure 2.**
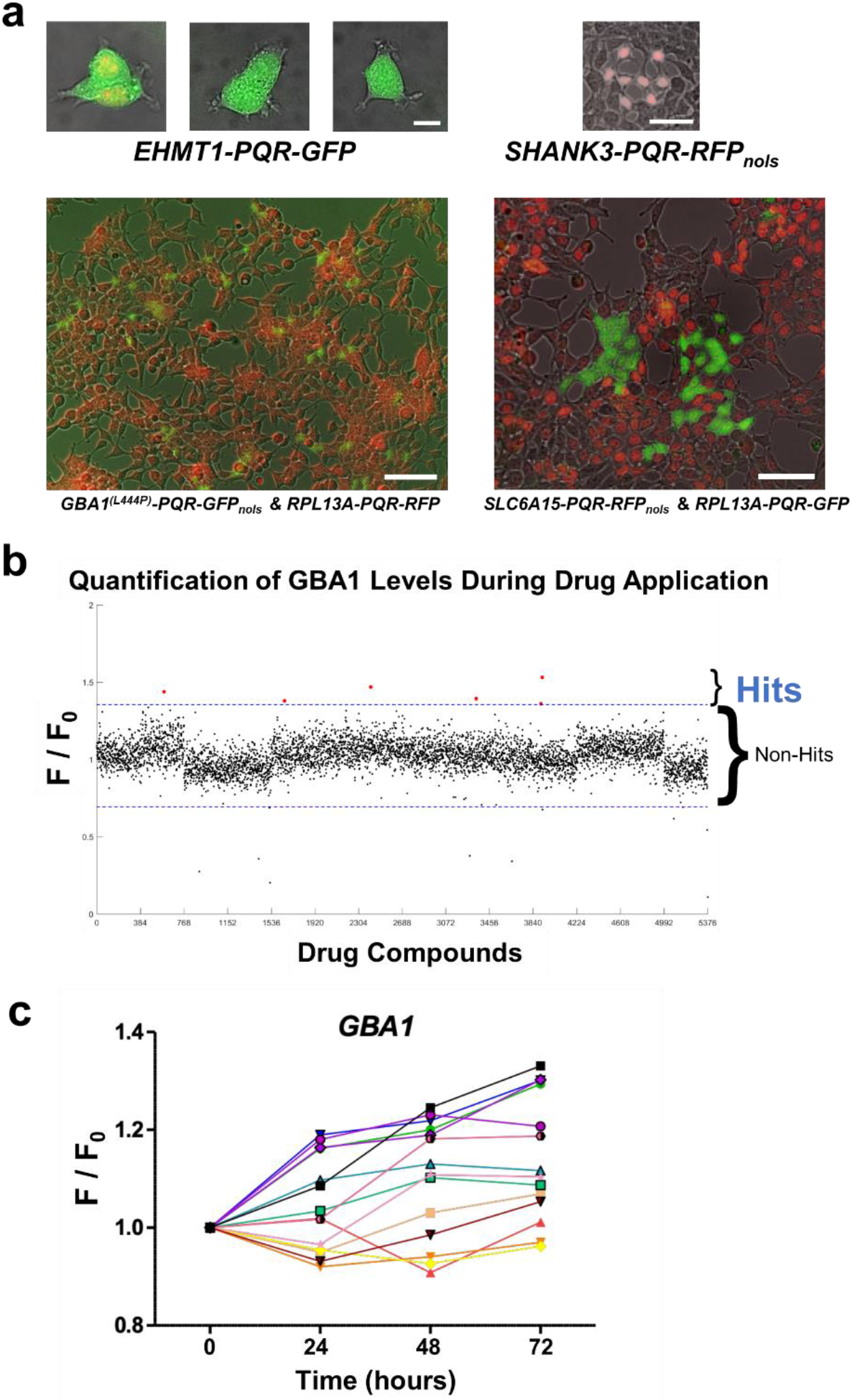
PQR can be used to screen drugs that increase endogenous protein levels. (a) Representative images of human HEK293T cells that have been genome-edited to insert a *PQR* and a fluorophore gene into four different target genes, *EHMT1, GBA1, SHANK3*, and *SLC6A15*. Scale bars are 10 µm (upper panels) and 50 µm (lower panels). (b) Representative example of a drug screen of >5,000 drug compounds using the *GBA1-PQR-GFP*; *RPL13A-PQR-RFP*_*nols*_ HEK293T cells at 24 hours after drug administration. Six drug compounds (red dots) significantly increased GBA1 levels normalized to RPL13A levels greater than three standard deviations (blue dotted lines). (c) Representative example of a validation screen of the drug hits (colored symbols/lines represent a single drug) on GBA1 protein expression over days.

Next, we screened through thousands of drug compounds using the *GBA1-PQR-GFP*; *RPL13A-PQR-RFP*_*nols*_ HEK293T cells, and the *SLC6A15-PQR-GFP*; *RPL13A-PQR-RFP*_*nols*_ HEK293T cells (**Figure 2b**). Cells were seeded at 2,000 cells per well in a 384-well plate and imaged using a high content imager with integrated liquid handling, and drug compounds were applied using an acoustic droplet liquid handler. The RPL13A-PQR control channel was monitored to exclude drugs that affected RPL13A as well as the target gene, and then the target gene PQR fluorescence values were normalized to the RPL13A levels. Other internal controls included randomly interspersed DMSO or media only controls in wells. However, artifacts near the edges of the 384-well plates were still sometimes observable, that the internal controls could not overcome. These “edge effects” are common in high-throughput screening (Mansoury et al., 2021) and are caused by a number of several factors, such as evaporation and temperature differences.

We then performed a validation screen of the top hits (**Figure 2c**), which were defined as those compounds that increased target protein expression greater than three standard deviations above the average signal at either 24, 48, or 72 hours. The top drug results from this follow-up screen can be verified *in vivo* using wildtype animals or disease models, or *in vitro* using human primary cells.

## Discussion

Traditional drug screening assays use cellular pathology or cellular phenotypes associated with the disease. Here, we show that drug screens for genetic disorders caused by abnormal protein expression can be performed using a *PQR* genetic tag and simple fluorescence microscopy. We used HEP-G2 and HEK293T cell lines as an easier model system for culturing, genome editing, and live imaging. We first confirmed the mRNA expression levels of the target gene within the cell line using proteinatlas.org (Ponten et al., 2011). However, high-throughput drug screens are not limited to cell lines, and can also be performed on cells differentiated from pluripotent stem cells. Using human induced pluripotent stem cells allows for drug screens in human neurons for neurological diseases and disorders. Screening for drugs using human neurons was previously not feasible as neurons are post-mitotic and cannot be cultured and passaged indefinitely, and thus the only means of a steady supply of neurons are pluripotent stem cells. We used induced pluripotent stem cells to insert a *PQR* into the target disease genes, *GBA1, SHANK3*, and *SLC6A15*. We used the *GBA1-PQR-GFP*_*nols*_ knock-in stem cells to differentiate these into dopaminergic neurons (**Figure 3**). These PQR knock-in neurons can not only be used for drug screening in a neurodegenerative disease-relevant cell type, but can also be combined with other cellular assays such as cell survival, metabolic activity, or morphology.

**Figure 3.**
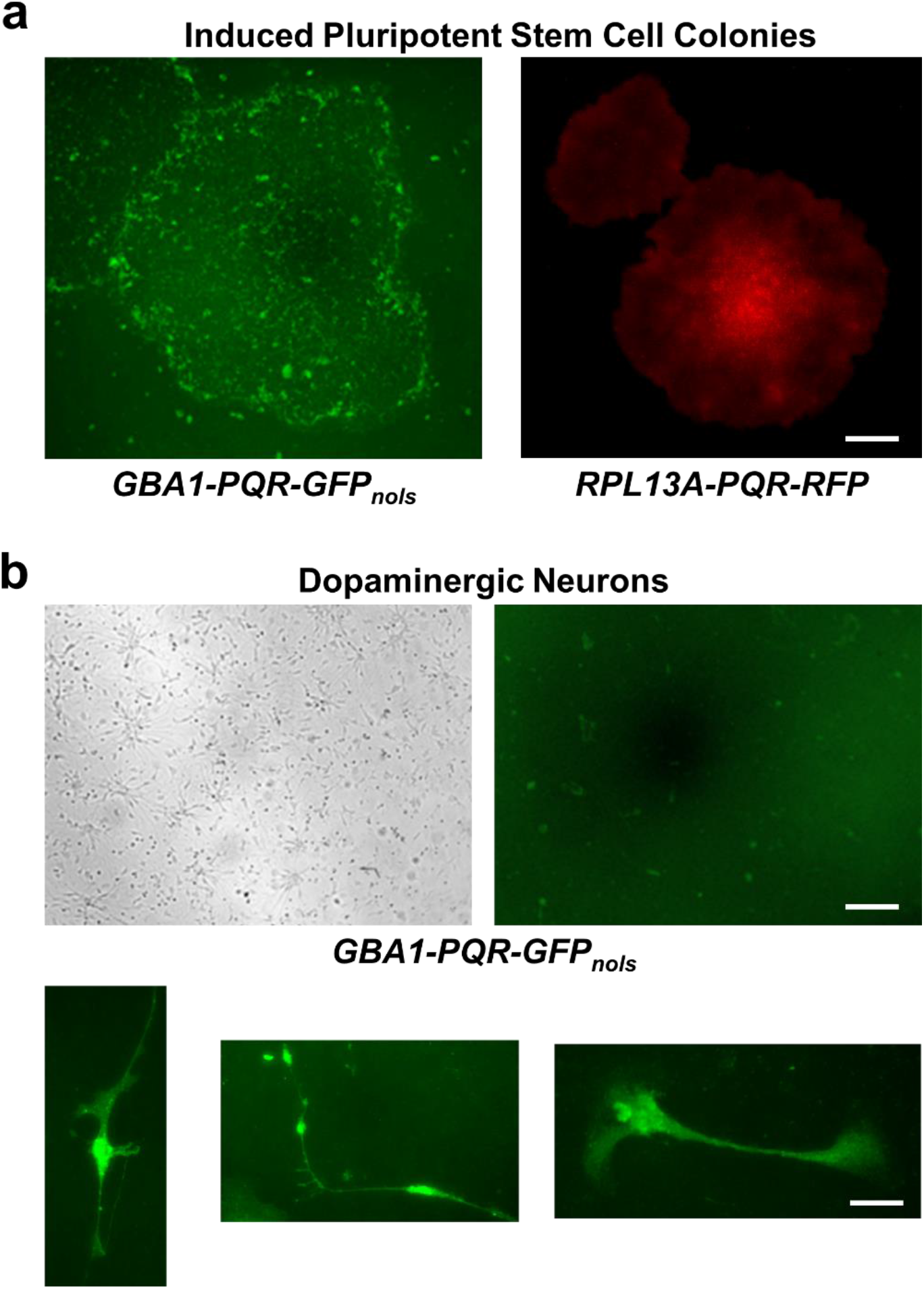
Human neurons can be used in drug screens for endogenous protein levels. (a) Representative examples of knock-in human induced pluripotent stem cell colonies with *GBA1-PQR-GFP*_*nols*_ (left image) or *RPL13A-PQR-RFP* (right image). Scale bar is 50µm. (b) Representative examples of *GBA1-PQR-GFP*_*nols*_ human dopaminergic neurons. Bottom images show single neurons stained for tyrosine hydroxylase in green. Scale bars are 50µm in top images, 10 µm in bottom images.

The protein expression drug screening approach has several disadvantages, however. The most critical disadvantage is the lack of knowledge of drug mechanism of action. A drug may change protein expression in a wide variety of ways, whether entering the cell or activating a surface receptor and then altering transcription of the target gene, or transcriptional regulation, or altering translation of the target protein, or translational regulation. Even if a drug lead has been validated *in vitro* and *in vivo* using multiple model systems and disease models, the mechanism by which it alters the target protein expression could remain a mystery. Without knowledge of how the drug acts, it will be difficult for medicinal chemistry to develop the drug further to increase the drug safety and efficacy, making it unattractive for pharmaceutical companies. One way to mitigate this disadvantage is to first determine whether the drug acts at the transcriptional level by using quantitative RT-PCR to measure changes in the mRNA in addition to changes in protein levels (Kays and Chen, 2019; Lo and Chen, 2019). Another mitigation can include screening with drug libraries of oligonucleotides, such as antisense oligonucleotides (or antisense therapy), since nucleotides have identifiable targets. Screening using FDA-approved drug libraries may circumvent this issue, since many drugs have unknown mechanisms of action, such as modafinil, metformin, and acetaminophen/paracetamol, as well as the majority of drugs that are prescribed off-label for unapproved indications, such as ketamine for treatment-resistant depression. As long as the drug lead is extensively validated pre-clinically, then the FDA-approved drug can potentially move directly into Phase 2 clinical trials to determine efficacy, and patients may be able to use the re-purposed drug off-label.

Another disadvantage of drug screening using PQR is that it cannot detect changes post-translationally, such as changes in protein turnover, protein modifications, or protein function. Additionally, for haploinsufficiency disorders if a gene mutation is a gain of function, where the mutant protein has deleterious effects on the cell, then a drug that increases the target protein would be detrimental because this would amplify the negative effects. In these cases, the target patient population would be those with loss of function mutations. Finally, some genes have very tightly regulated protein expression where too little or too much can result in abnormal cellular function. *SHANK3* is one such gene, where complete loss of one copy of the gene results in Phelan–McDermid syndrome (Monteiro and Feng, 2017; Yi et al., 2016), and too much expression of SHANK3 causes manic-like behaviors (Han et al., 2013). Thus, any drug that alters protein abundance will require detailed dosage, pharmacodynamic, efficacy, and safety characterizations.

There are several advantages to drug screening using protein levels. Using PQR, time-lapse imaging can be performed in single cells to measure protein expression before, during, and after drug application (**Figure 2c**) to measure the pharmacodynamics of the drug. The PQR approach measures protein amounts, which is a more accurate reflection of the actors within a cell, rather than mRNA levels. Thus, drug-induced changes that occur post-transcription, such as protein synthesis regulation, can also be detected. The fluorophore used with the *PQR* tag can also offer several advantages, such as the use of fluorescent timer proteins and photoconvertible proteins, such as Dendra (**Figure 1**). Fluorescent timer proteins change their emission from blue to red over a period of hours (Subach et al., 2009; Subach et al., 2022). Thus, the rate of protein synthesis can be examined by making repeated measurements using the blue channel, and this can be performed regularly over time scales of days of drug application. The photoconvertible fluorescent protein Dendra normally emits green fluorescence, but can be permanently photoconverted to emit red fluorescence by UV illumination (Chudakov et al., 2007). The rate of protein synthesis can be measured at any arbitrary time by UV illuminating the cell to convert the existing signal to the red channel, and new protein synthesis will then occur within the green channel. Thus, fluorescent timer proteins and photoconvertible fluorescent proteins shift accumulated fluorescence into a different channel, so that new protein synthesis events can be repeatedly measured. Finally, other advantages of using PQR to measure protein levels for drug screening is the ability to use patient-derived cells, including induced pluripotent stem cells (**Figure 3**). The genotype of these cells can be changed using CRISPR-Cas9 to re-create (**Figure 2a**) or correct patient mutations in the same genome editing step as the *PQR* insertion.

Further validation of drug hits can include *in vivo* mouse models. Conceptually, the same protein quantification using PQR during drug application can be performed in a knock-in mouse. Drug candidates need to be optimized to improve pharmacokinetic and pharmacodynamic characteristics such as absorption, distribution, metabolism, and toxicity. Ideally, in each round of drug development the drug lead is continually re-validated using rapid *in vitro* and *in vivo* assays. Thus, a PQR knock-in mouse similar to the PQR knock-in human cells would be a useful holistic and whole-animal assay for several reasons. First, tracking the endogenous protein expression of a target disease gene over time will itself be an important model for the disease. Second, imaging how the target protein expression changes within different tissues is critical for pharmacokinetics as the target protein expression may run down over time with drug duration, dosage, route, or circadian cycle. Third, imaging the target protein expression spatially across different tissues will aid in the pharmacodynamic evaluation of drug efficacy. Fourth, imaging target protein expression over time across different brain regions is important for models of neurological diseases. Fifth, imaging the target protein expression can be combined with mouse models of the disease, by crossing the mice or using chemotoxic agent models. Sixth, using near-infrared fluorescent proteins allow for nanometer to centimeter scale *in vivo* imaging (i.e., fluorescence microscopy to whole animal imaging) (Matlashov et al., 2020; Oliinyk et al., 2022). This type of PQR knock-in mouse will allow for whole animal imaging through the skin, before, during, and after drug administration to track the target protein expression throughout the body in real time. Imaging these mice through the skin, combined with drug tracing through nuclear medicine imaging, would allow for much more accurate, rapid, and cheaper pharmacology and toxicology measurements while still monitoring drug efficacy simultaneously.

### Experimental Procedures

#### Cell Culture

HEP-G2 and HEK293 cells were cultured at 37°C under 5% CO2 in Dulbecco’s Modified Eagle Medium (Wisent, St-Bruno, QC). All media were supplemented with 10% fetal bovine serum (FBS) (Wisent), and 100 units/mL penicillin (Life Technologies, Carlsbad, CA) and 100 μg/mL streptomycin (Life Technologies). iPSCs were cultured in growth media mTESR plus, or mTESR1. The monolayer neural induction protocol (STEMCELL Technologies) was used to obtain NPCs. Briefly, iPSCs were seeded at 3×10^6^ cells / mL on 6-well plates coated with Matrigel (Corning) containing Neural induction media with SMADi supplemented with 10 µM Y-27632 (STEMCELL Technologies) on day 0. From day 1 to day 6, media changes were performed with Neural induction media and SMADi alone. Cells were passaged after at least 6 days for 3 rounds.

To obtain dopaminergic neurons, we used the STEM™ differentiation and maturation kit (STEMCELL technologies). Briefly, NPCs were plated onto 6-well plates coated with PLO/laminin and resuspended in 2 mL dopaminergic differentiation media. Media change was performed every day until cells reached 90% confluency. Cells were passaged and seeded at a density of 6×10^4^ cells / mL and resuspended in dopaminergic maturation media 1 (STEMCELL Technologies) for 4 days. Media changes were performed every day. On day 5, we changed to Dopaminergic Media 2 (STEMCELL Technologies) for another 5 days.

#### CRISPR/Cas9 Genome Editing

The genomic sequences of each of the target genes, *Apolipoprotein E* (*APOE*), *Euchromatic Histone-lysine N-methyltransferase 1* (*EHMT1*), *β-Glucocerebrosidase* (*GBA1*), *SH3 and multiple ankyrin repeat domains 3* (*SHANK3*), and *Sodium-dependent neutral amino acid transporter B(0)AT2* (*SLC6A15*), and *Ribosomal protein L13A* (*RPL13A*) were PCR amplified from the human cell lines. The *Protein Quantitation Reporters* (*PQRs*) used were as described for mammalian genomes (Lo et al., 2015). The target gene fragments with the *PQR* and fluorescent protein gene were assembled into cassettes and placed between the homology arms using In-Fusion Cloning (TAKARA Bio). The homology arms did not include the endogenous promoter, thus preventing the expression of the transgene until the in-frame genomic integration at the correct locus occurred during genome editing. Sizes of homology arms were ∼1 kb.

Four different single guide RNAs (sgRNAs) were designed to guide the Cas9 nuclease to make a double strand break at the end of the coding region of each gene. They were designed in a 20 bp DNA oligonucleotide format. Individual sgRNAs were cloned into a dual promoter plasmid, such that both the sgRNA and Cas9 were expressed from the same vector. Transfection was achieved using Lipofectamine 3000 (ThermoFisher) or electroporation using a NEPA21 (NepaGene). 800 ng of *CRISPR-Cas9* plasmid DNA were co-transfected with 800 ng of repair template circular plasmid in 12-well plates. Efficiency of sgRNA/Cas9 was evaluated five days after transfection by cellular fluorescence and genotyping.

Genotyping experiments were performed in experimental duplicate. Integration of *PQR* into the endogenous genomic locus was validated by genomic DNA extraction six days post-transfection and genotyping using primers outside of and within the homology arms of the repair template. The 5’ (upstream) and 3’ (downstream) ends of the insertion were probed with two sets of primers and the endogenous locus was PCR amplified. Restriction digests were then performed on PCR products at sites specific for *PQR*. All genomes were sequenced to identify the *PQR* and genomic junctions.

#### Drug Screening

T0901317 (Abcam) was dissolved in DMSO and applied at 250nM to 10µM concentrations. Fluorescence images of >100 cells/well in 384-well plates were taken using a Perkin-Elmer Opera Phenix High Content Imager. Liquid handling was performed using a Beckman Biomek Integrated Robotic System, and drug compounds were applied using a Labcyte Echo 555 Acoustic Droplet liquid handler.

#### Image Analysis and Statistical Analysis

Images were analyzed using the Perkin-Elmer Opera Harmony software or ImageJ, and selected on the basis of identification of single cells with normal morphology. Images were adjusted for contrast and brightness only. Average pixel intensities were calculated for each well. We verified that the RPL13A-PQR control channel fluorescence did not increase or decrease significantly throughout the time-lapse imaging. Student’s *t*-tests were performed to test the null hypothesis that the average fluorescence values from the different conditions were the same, from two-tailed distributions of similar variances (*p* < 0.05).

## Author Contributions

B.E.C. designed the experiments and supervised the project. F.E., I.K., C.L., Y.L., and B.E.C. performed experiments and analyzed the data. F.E. and B.E.C. wrote the manuscript.

## Acknowledgments

The authors thank Shruti Sharma and Uyen Do for assistance with experiments. This work was supported by grants (to B.E.C.) from the Canadian Institutes of Health Research (148882), GeneSpark and the International Kleefstra Syndrome Foundation, the Sandra and Alain Bouchard Foundation, the Weston Brain Institute, and the Consortium Québécois sur la Découverte du Médicament.

## Competing Interest Statement

B.E.C. is an inventor on a patent on PQR. B.E.C., I.K., and C-A. L. are founders of a company based on the work in this manuscript.

Correspondence and requests for materials should be addressed to

